# Retina-specific targeting of pericytes reveals structural diversity and enables control of capillary blood flow

**DOI:** 10.1101/2020.05.29.124586

**Authors:** Elena Ivanova, Carlo Corona, Cyril G. Eleftheriou, Paola Bianchimano, Botir T. Sagdullaev

**Affiliations:** Burke Neurological Institute, White Plains, NY 10605; Department of Ophthalmology, Weill Cornell Medicine, New York 10065

**Keywords:** Mural cells, pericyte, precapillary sphincter, retina, optogenetics, Halorhodopsin

## Abstract

Pericytes are a unique class of mural cells essential for angiogenesis, maintenance of the vasculature and are key players in microvascular pathology. However, their diversity and specific roles are poorly understood, limiting our insight into vascular physiology and the ability to develop effective therapies. Here, in the mouse retina, a tractable model of the CNS, we evaluated distinct classes of mural cells along the vascular tree for both structural characterization and physiological manipulation of blood flow. To accomplish this, we first tested three inducible mural cell-specific mouse lines using a sensitive Ai14 reporter and tamoxifen application either by a systemic injection, or by local administration in the form of eye drops. Across three lines, under either the PDGFRb or NG2 promoter, the specificity and pattern of Cre activation varied significantly. In particular, a mouse line with Cre under the NG2 promoter resulted in sparse TdTomato labeling of mural cells, allowing for an unambiguous characterization of anatomical features of individual sphincter cells and capillary pericytes. Furthermore, in one PDGFRb line, we found that focal eye drop application of tamoxifen led to an exclusive Cre-activation in pericytes, without affecting arterial mural cells. We then used this approach to boost capillary blood flow by selective expression of Halorhodopsin, a highly precise hyperpolarizing optogenetic actuator. The ability to exclusively target capillary pericytes may prove a precise and potentially powerful tool to treat microcirculation deficit, a common pathology in numerous diseases.

## Introduction

Pericytes comprise a heterogeneous population of mural cells with a variety of functions, yet their identity remains a topic of intense debate [1]. In the CNS, capillary pericytes have the highest densities and ratio to endothelial cells, and are thought to be essential to angiogenesis, neovascularization, blood-brain-barrier function and dynamic control of blood flow [2]. Accumulating evidence links pericyte failure to major brain pathologies such as stroke, Alzheimer’s disease, BBB impairment, neurodegeneration and traumatic brain injury [3,4]. Therefore, a better understanding of pericyte subtypes and their unique roles is needed for the development of effective therapeutic strategies.

Several inducible Cre mouse lines have been developed specifically to dissect pericyte contribution to CNS physiology and pathology, expressing Cre under either the PDGRFb (Table 1, line 1-2) or the NG2 promoters (Table 1, line 3). The PDGRFb-mT/mG line [5], marks a majority of mural cells across the CNS at P6 [6]. Conversely, the CSPG4-mT/mG line, assessed in the same study demonstrated minimal expression in mural cells as well as off-target expression in glial cells. However, these lines were not validated in adulthood with no display of off-target expression and no evaluation of the specificity for selective mural cell types. The PDGFRb-cre mouse was validated in the CNS, including the retina at a variety of developmental stages using two reporter lines [7]. When crossed with a sensitive TdTomato reporter [8] around 84% of mural cells were positive at P5. At 6 weeks, all types of mural cells were found to express the reporter. Therefore, lack of experimental consistency makes it hard to fully utilize this approach to pericyte physiology. Moreover, no organ specific targeting was established, which is important to the development of therapeutics.

**Table 1.**
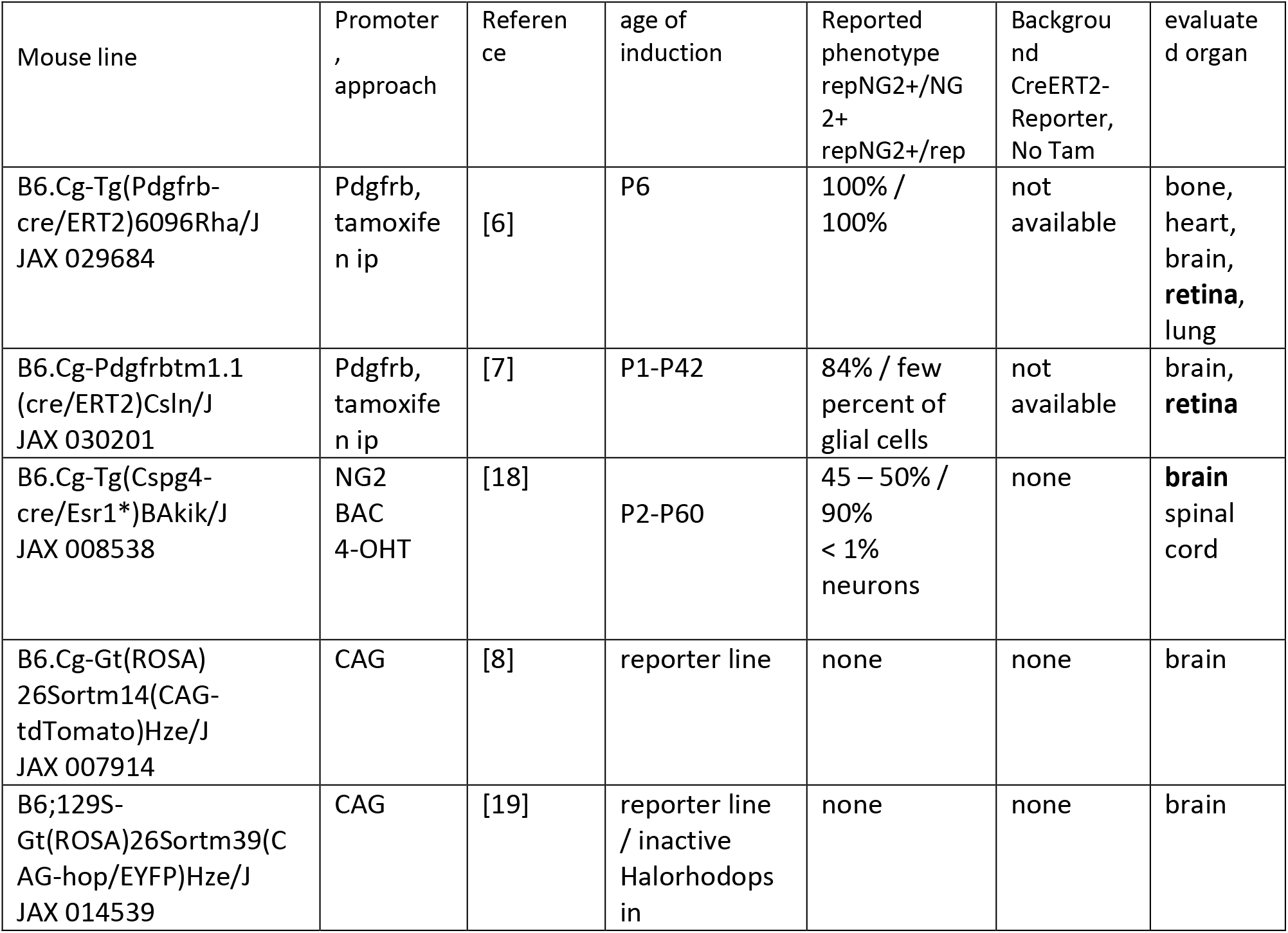
Commercially-available mouse lines with tamoxifen-inducible Cre in vascular mural cells.

To address these limitations in the characterization of retinal mural cells, we characterized activation patterns of tamoxifen-inducible mural cell specific CreERT2 mouse lines with a highly sensitive Ai14 TdTomato reporter line. The three mouse lines described above (Table 1, lines 1-3) drive expression of TdTomato under either the NG2 or PDGRFb promoter, enabling in vivo, ex vivo and structural characterization in the retina. Indeed, the retina is an accessible layered part of the brain that has been used as an ideal model to study CNS angiogenesis [9] and blood flow [10] in health and pathological conditions. Angiogenesis and vascular maintenance are essential for proper organ development and function, particularly in the highly metabolically demanding central nervous system [11]. In contrast to the rest of the brain, the retinal vascular system develops after birth and is very well-characterized spatially and temporally [9]. All types of blood vessels are present in the retina. The architecture of the retina and its vascular system is conserved among the vertebrates [12]. Direct non-invasive assessment of developing vasculature at cellular level is possible using advanced confocal and two-photon microscopy. Additionally, the retina is the most metabolically demanding tissue in the human body by weight [13], and it must therefore be nourished by a large blood supply. There are a number of diseases in the retina that are affected or driven by pericyte pathology [14].

Using sparse Cre-activation in isolated mural cells, we characterized the morphology of mural cells along distinct regions of the vascular tree. Specifically, the structure of sphincter pericytes and capillary pericytes was determined. With an eye-drop tamoxifen treatment we were able to selectively activate Cre in capillary pericytes but not in arterial smooth muscle cells. As a proof of concept for pericyte-driven optogenetic therapeutics, we selectively expressed Halorhodopsin in capillary pericytes and achieved Halorhodopsin-assisted blood flow enhancement in retinal capillaries *in vivo*.

## Methods

### Animals

All animal procedures were performed in accordance with the Institutional Animal Care and Use Committee of Weill Cornell Medicine, and in accordance with the National Institutes of Health Guide for the care and use of laboratory animals. All mouse lines were purchased from Jackson Laboratories (Table 1).

To evaluate Cre activation patterns, CreERT2 mouse lines were crossed with either Ai14 or Halorhodopsin reporter lines. All experiments were conducted in mice heterozygous for both Cre and reporter genes. After genotyping, one group of animals was treated with standard intraperitoneal (ip) injections of 100 ul 20 mg/ml tamoxifen in corn oil daily for five consecutive days (Jackson Laboratories protocol). The other group was treated by 5 ul 5 mg/ml tamoxifen in corn oil three times a day for five consecutive days [15]. As controls, Cre and TdTomato-positive mice were treated with corn oil either via ip or via eye drops. Three to ten days after the final tamoxifen treatment, tissue was collected and analyzed for efficiency and specificity of TdTomato expression.

### Two-photon recordings of retinal capillary blood flow

For in vivo blood flow characterization, the tail vein was injected with 200 μl Sulforhodamine 101 (SR101, 10 mM, Sigma, #S7635) 10 min prior to anesthesia. Each animal was initially anesthetized with a mixture of 75 mg/kg ketamine and 7.5 mg/kg xylazine. The pupils were dilated with a 0.5% tropicamide ophthalmic solution, and a coverslip was placed on each eye with GONAK ophthalmic solution. Mice were mounted with a SG-4N mouse head holder (Narishige) on an upright ThorLabs Bergamo II two-photon microscope. During recordings, mice were placed on a heat pad and covered with a blanket, to maintain physiological temperature. Blood flow was measured using a 10X super apochromatic objective with a 7.77 mm working distance and 0.5 NA (TL10X-2P, ThorLabs, Newton, NJ). Using a long working distance, we first placed the objective as close to the eye as possible with the focus beyond the retina and without any fluorescent illumination. The focus was achieved in two-photon mode by moving the objective away from the eye. SR101 was illuminated with a 920 nm wavelength and the measurements were taken at a rate of 116 frames per second. At first, baseline blood flow was recorded for 30 s. During this time, superficial, intermediate and deep capillary layers were measured by manually focusing through the retina; each vascular layer was recorded for several seconds. Following baseline recordings, the eye was illuminated for 20 s through the same objective using a 550-575 nm red LED. After light stimulation, each vascular layer was recorded for a total of 30 s. The time-lapse images were corrected for background, filtered with a Gaussian filter, and, when necessary, stabilized in Fiji (NIH). A stabilization plugin was used to eliminate strong movements of capillaries during deep animal breathing. Blood flow was estimated as the number of blood cells passing through a capillary per second. This analysis was performed in Fiji by plotting fluorescence profiles across blood vessels for every frame. In the resulting plot, troughs indicate the passage of a blood cell, which was darker relative to the plasma labeled with SR101. The peaks were automatically detected in Microsoft Excel and verified by visual inspection of the original data [16].

### Immunocytochemistry

As described previously [17], following euthanasia, the eyes were removed and placed in bicarbonate-buffered Ames’ medium (Sigma Aldrich, St Louis, MO) equilibrated with 95% O2 and 5% CO2. The cornea was removed by an encircling cut above the ora serrata, and the iris, lens, and vitreous were extracted. The remaining eyecup, with the retina still attached to the pigment epithelium was submersion-fixed on a shaker in freshly prepared 4% paraformaldehyde in 0.1 M phosphate saline (PBS, pH = 7.3) for 20 min at room temperature. After fixation, the eye cups were washed in PBS for 2 h and the retinas were detached from the eye-cups. Isolated retinas were blocked for 10 h in a PBS solution containing 5% Chemiblocker (membrane-blocking agent, Chemicon), 0.5% Triton X-100, and 0.05% sodium azide (Sigma). Primary antibodies were diluted in the same solution and applied for 72 h, followed by incubation for 48 h in the appropriate secondary antibody. In multi-labeling experiments, tissue was incubated in a mixture of primary antibodies, followed by a mixture of secondary antibodies. All steps were completed at room temperature. After staining, the tissue was flat-mounted on a slide, ganglion cell layer up, and coverslipped using Vectashield mounting medium (H-1000, Vector Laboratories). The coverslip was sealed in place with nail polish. To avoid extensive squeezing and damage to the retina, small pieces of a broken glass cover slip (Number 1 size) were placed in the space between the slide and the coverslip. Retinal wholemounts were imaged under a Leica SP8 confocal microscope using 20x air and 63x oil objectives.

The following primary antibody were used: mouse anti-SMA (Sigma, A5228, 1:2000, RRID:AB_262054), goat anti-CD31/PECAM-1 (RD, AF3628, 1:6000, RRID:AB_2161028), mouse anti-glutathione synthetase (MilliporeSigma, MAB302, 1:2000, RRID:AB_2110656), goat anti-PDGFRb (RD, AF1042, 1:8000, RRID:AB_2162633), and rabbit anti-Cre recombinase (Cell Signaling, Cat. 15036, 1:2000, RRID:AB_2798694). We also used *Griffonia simplicifolia* isolectin (GS-IB4 Alexa Fluor 488, I21411, RRID:AB_2314665). Isolectin stock solution was made using 500 μg isolectin powder dissolved in 500 μl PBS-calcium solution: 8.1 mM Na2HPO4 dibasic, 1.9 mM NaH2PO4 monobasic, 154 mM NaCl, 1 mM CaCl2. Isolectin stock solution was diluted 1:400 and applied with the secondary antibodies.

### Statistical analysis

Statistical analysis was performed in Microsoft Excel using t-test. The data are presented as mean ± standard deviation, unless otherwise indicated. The number of samples (n) indicates the number of samples per group.

## Results

### Evaluation of inducible mural cell-specific Cre mouse lines for use in the retina

We used commercially available mouse lines with tamoxifen-inducible Cre in vascular mural cells (Table 1): B6.Cg-Tg(Pdgfrb-cre/ERT2)6096Rha/J (shortly, Pdgfrb-CreRha), B6.Cg-Pdgfrbtm1.1 (cre/ERT2)Csln/J (shortly, Pdgfrb-CreCsln), and B6.Cg-Tg(Cspg4-cre/Esr1*)BAkik/J (shortly, Cspg4-Cre).

These were crossed with bright Ai14 TdTomato reporter line (abbreviated Tom, [8]) to reveal Cre expression and activation. Offspring heterozygous for Cre and TdTomato were treated with tamoxifen starting at age P22 (Figure 1). One group of animals was treated with standard intraperitoneal injections (ip) of 100 ul 20 mg/ml tamoxifen in corn oil daily for five consecutive days (Jackson laboratories protocol). The other group was treated with 5 ul eye drops of 5 mg/ml tamoxifen in corn oil three times a day for five consecutive days [15]. As controls, Cre-TdTomato-positive mice were treated with corn oil either via ip or via eye drops. In three to ten days after the last tamoxifen treatment, the tissue was collected and analyzed for efficiency and specificity of TdTomato expression (Figure 1b-i). We focused on the superficial layer as it contains all types of the blood vessels: artery (positive for smooth muscle cells), relay (a part of the vascular tree, originating in capillary and extending toward feeding artery [20]), capillary, and vein (Figure 1c,d). Morphologically unique mural cells (magenta text) could be readily identified along the distinct vascular tree segments (white text), indicative of their function-specific vascular residence. Individual TdTomato-positive cells were further distinguished in high resolution (Figure 1e-i) using established functional markers for smooth muscle actin (SMA), endothelial cells (CD31) and basement membrane (isolectin) [16]. Thus, a nuanced and type-specific approach is required to evaluate the roles of diverse mural cell types on the vascular tree and to control them in genetic models.

**Figure 1.**
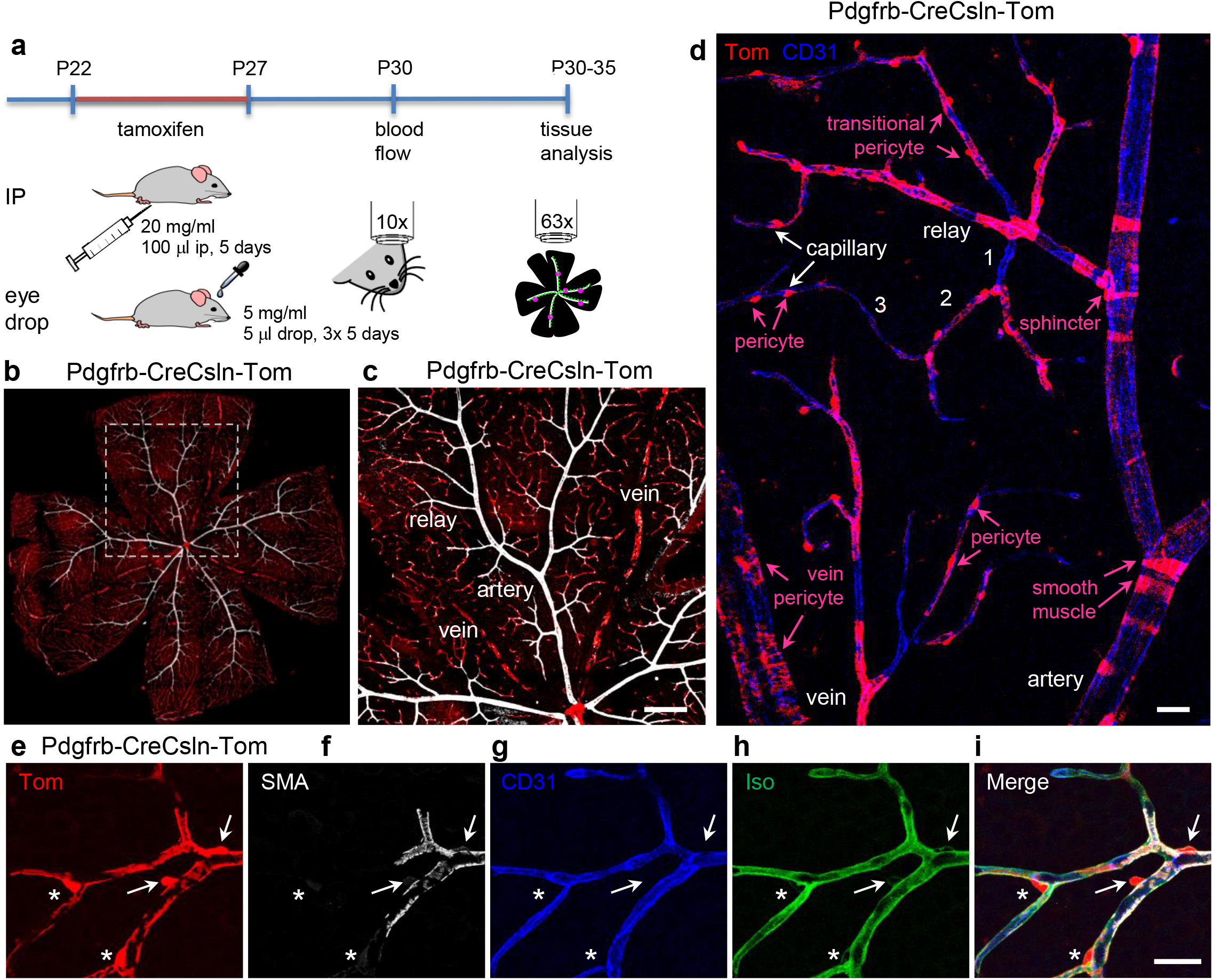
Experimental approach for activation of CreERT2 in retinal vascular contractile cells and their assessment with a TdTomato reporter. (a) Tamoxifen was delivered either by intraperitoneal injection or via eye drops. (b) Evaluation of retinal wholemounts after tamoxifen-mediated activation of CreERT2. Retinas were labeled for smooth muscle actin (SMA, white) to identify arteries, red fluorescence from TdTomato reporter line indicates active Cre. B6.Cg-Pdgfrbtm1.1 (cre/ERT2)Csln/J mouse is shown. Scale bar 200 um. (c) High magnification image from highlighted area in (b). Scale bar 200 um. (d) Hierarchy of the vascular tree and diversity of mural cells along the vascular branches. Vascular tree was visualized by immunolabeling for CD31, a marker of endothelial cells. Numbers indicate capillary order. Scale bar 25 um. (e-i) Identification of TdTomato-positive cells using markers for SMA-positive contractile cells (SMA, white), endothelial cells (CD31, blue), basement membrane which covers pericytes and SMA cells (Iso, green). TdTomato-positive cells were identified as smooth muscle cells (arrows) and pericytes (asterisks). Scale bar 25 um.

### Identification of morphological polarization of mural cells using sparse labeling: pericytes and sphincter

To examine diverse mural cells, we used the Cspg4-Cre mouse line that produces sparse labeling of diverse mural cells [18]. When crossed to the TdTomato reporter line, the resulting Cspg4-Cre-Tom mouse had no “leaky” expression of TdTomato in the absence of tamoxifen (Figure 2a). When the mouse was ip injected with tamoxifen, a small number of mural cells became labeled (Figure 2b). In addition to brightly labeled mural cells, a relatively weak labeling was also found outside of the vasculature (Figure 2c). Similarly, in the reporter mouse alone (Figure 2d), the same structures were weakly labeled (Figure 2e). These weakly positive structures were double-labeled for glutamine synthetase, a marker for Muller cells (Figure 2f).

**Figure 2.**
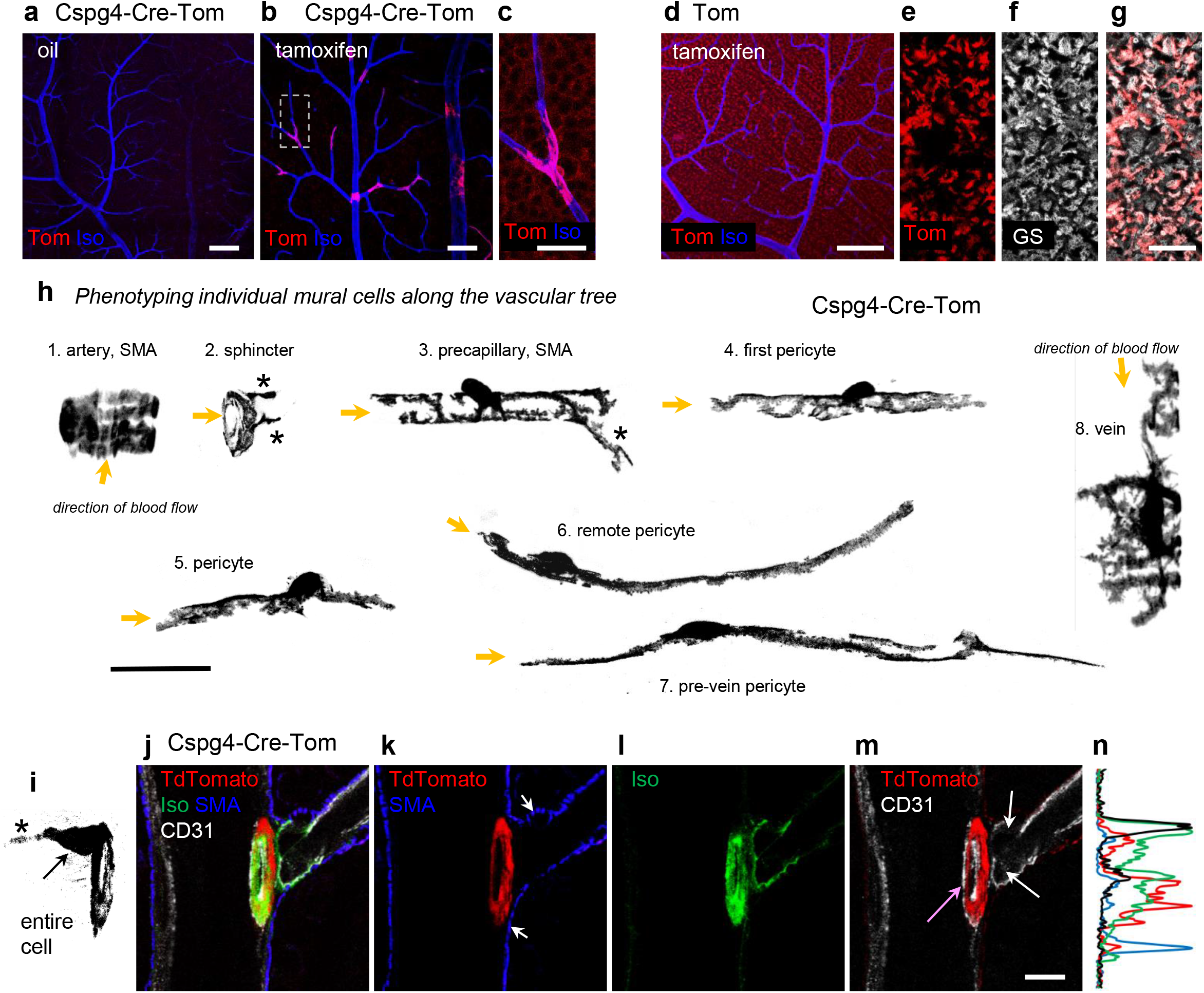
In Cspg4-Cre line, tamoxifen treatment results in Cre-induction across a sparse population of contractile vascular cells. (a) In the oil injected mice, no “leaky” activation of CreERT2 was detected. Scale bar 100 um. (b) Tamoxifen ip induced TdTomato expression in sparsely distributed vascular mural cells. Scale bar 100 um. (c) High resolution image from the area highlighted in (b). When TdTomato brightness was strongly amplified, weak TdTomato signal was detected around soma in the ganglion cell layer. Scale bar 20 um. (d) In the reporter mouse, weak TdTomato signal was found in the absence of Cre. Please note that the TdTomato signal was amplified in comparison with settings for (a-b). Scale bar 100 um. (e-g) High magnification of an area from (d) taken at the inner limiting membrane, where red TdTomato signal was colocalized with Muller cell marker, glutamine synthetase (GS, white). Scale bar 20 um. (h) Phenotypic characterization of sparely labeled by TdTomato mural cells along the vascular tree. Cells expressing SMA are marked. Asterisks mark fine processing along branching vessels. Scale bar 25 um. (i-n) Structural assessment of the artery branching point. Scale bar 10 um. (i) A projection through the sphincter cell. (j) The branching complex labeled for endothelial cells (CD31, white), smooth muscle actin (SMA, blue), sphincter (TdTomato, red), and basement membrane (Iso, green). (k) Sphincter cell (red ring) has very low SMA level in comparison with surrounding SMA-positive cells (blue labeling, arrowheads). (l) The branching point is strongly insulated by especially thick basement membrane (green). (m) Two endothelial cells are located at the base of the branch (CD31, white) with additional endothelial cells surrounding the entrance into the branch (pink arrow). All endothelial cells described here were also well insulated with the basement membrane (l).

The majority of Muller cells were double labeled (Figure 2g) and no labeling of any mural cells was found in the reporter line. The Cspg4-Cre-Tom mice treated by tamoxifen eye drops had sparse TdTomato expression in the mural cells, with the density lower than that exhibited by ip-treated mice.

Next, in the Cspg4-Cre-Tom mouse we characterized all types of positive mural cells (Figure 2h-n). Starting from the artery, we imaged progressively along the vascular tree towards the vein (Figure 2h). The SMA-positive cells on the artery had a compact and symmetric appearance with wide ribbon-like processes wrapping the artery from below. The “sphincter” cell positioned at the transition between the artery and precapillary region [21] completely encircled the branch opening with fine processes expanding onto the downstream vessel. Precapillary SMA-positive mural cells had symmetric shapes, occasionally with a fine process onto the neighboring branch (Figure 2h, asterisk). Pericytes had no SMA expression and had asymmetric processes with respect to their cell body. In the direction towards the feeding artery, processes were short and dense, wrapping the capillary lumen, and resembling those seen in precapillary mural cells. The processes towards the downstream direction were long and straight. Pericytes on finest remote capillaries had the longest processes. In contrast, the pre-vein capillary often had denser processes at the vein site. The vein pericytes had a “spider” like appearance with a dense network of fine processes.

Using the Cspg4-Cre-Tom mouse, we evaluated the sphincter cells (Figure 2i-n), a unique structure that is believed to play a role in gating blood inflow into the arterial branch [21–23]. The soma of these cells was commonly located on the major artery, although a few cells had their somas on the branch. A fine process (asterisk) protruded from the body and was also positioned on the main artery. The major processes of this cell tightly and continuously enwrapped the entrance to the branching vessel, creating a diaphragm. Interestingly, the sphincter cells had much lower expression of SMA relative to neighboring mural cells (Figure 2k, arrows). It was also insulated from nearby cells by a thick basement membrane (Figure 2l). This insulation could be protective against shear stress. It could also electrically decouple and chemically isolate the sphincter from the rest of the branch [22,24]. Typically, three endothelial cells surrounded the sphincter, with two at the entrance to the branch and one covering the artery side (Figure 2m). Figure 2n shows a profile through the sphincter and surrounding elements, where the basement membrane strongly insulates the complex (Isolectin Alexa488, green) and SMA has a relatively low expression in the sphincter.

Thus, the Cspg4-Cre mouse line can be used to sparsely label or control mural cells. In this line, the mural cells were randomly induced, and the majority of mural cells could be found without preference for any particular type. This enabled a thorough characterization of previously unknown mural cell subtypes. anatomy of uncommon types of mural cells could be examined.

### Ubiquitous and pericyte-specific targeting of mural cells in the retina

To achieve wide-spread targeting of mural cells, we used the Pdgfrb-CreRha and Pdgfrb-CreCsln mouse lines (Figure 3). In the absence of tamoxifen, the retinas of Pdgfrb-CreCsln-Tom mice exhibited only trace labeling of pericytes and no SMA cell labeling (Figure 3a). With tamoxifen ip injections, robust expression was achieved in all mural cells (Figure 3b and insert). As shown in the TdTomato reporter line (Figure 2d and e), weak labeling was also observed in many Muller cells. Interestingly, eye drop-tamoxifen application produced selective and complete labeling of pericytes and transitional cells on precapillary regions, but not of SMA cells on main arteries (Figure 3c).

**Figure 3.**
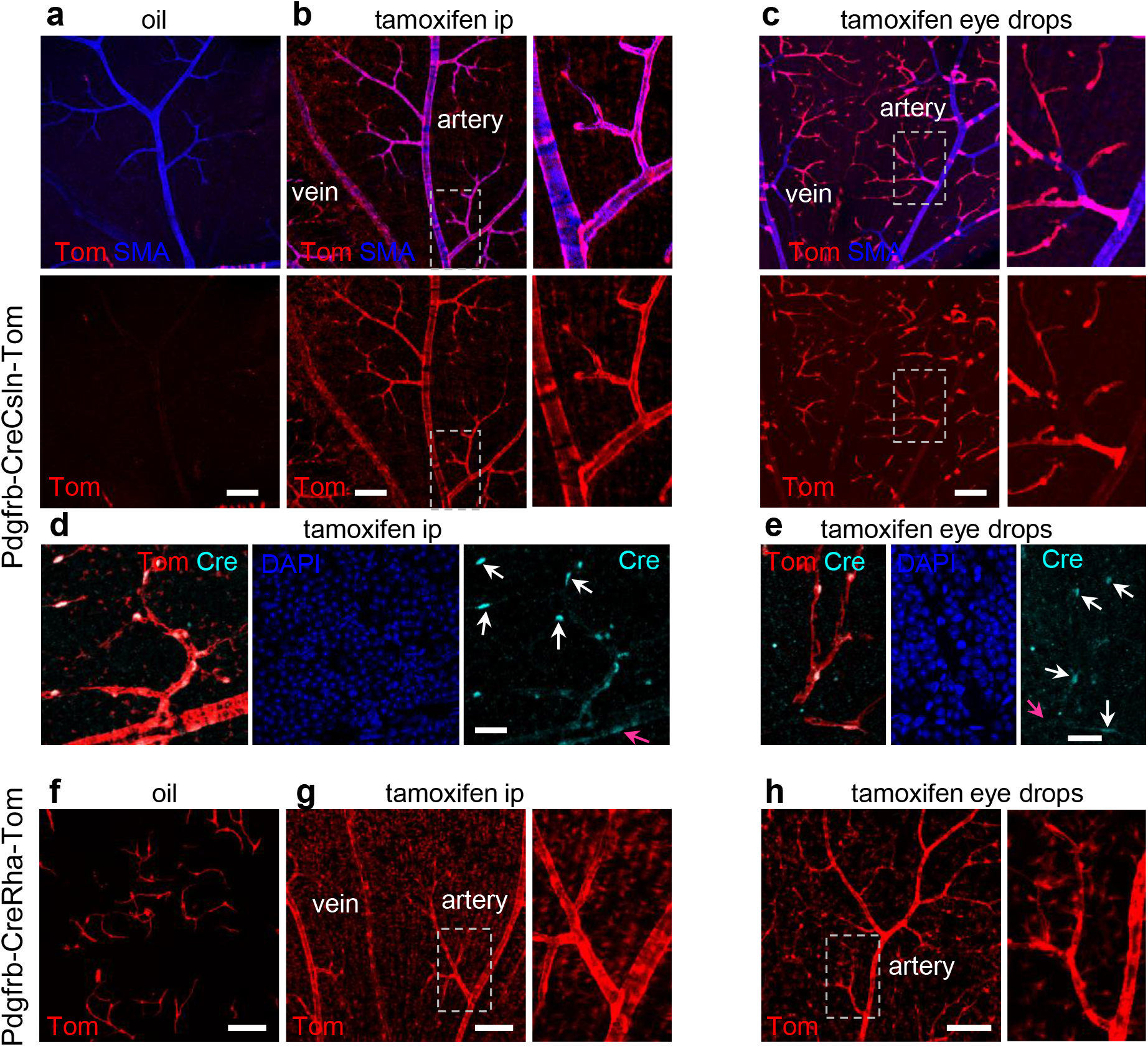
Comparison of two Pdgfrβ-CreERT2 lines for baseline activation of Cre and specificity of Cre activation after tamoxifen induction. (a) Pdgfrβ-CreCsln-Tom line had a very few TdTomato-positive mural cells without tamoxifen induction. (b) After ip tamoxifen treatment, Pdgfrβ-CreCsln-Tom line exhibited prominent TdTomato-labeling in all types of mural cells. All mural cells were positive. Weak TdTomato-expression was detected in Muller cells (magnified region). (c) In Pdgfrβ-CreCsln-Tom treated with tamoxifen eye drops, TdTomato was found predominantly in the capillaries and significantly less frequent in SMA cells and mural cells on veins. (d) After ip tamoxifen treatment, Pdgfrβ-CreCsln-Tom line showed Cre immunoreactivity in mural cell nuclei. The intensity of Cre immunolabeling was higher in pericytes (white arrows) and less in SMA cells (magenta arrows). (e) After tamoxifen eye drop treatment, Pdgfrβ-CreCsln-Tom line showed weak Cre immunoreactivity in pericytes (white arrows). Cre immunolabeling in SMA cells was below detection threshold (magenta arrow). (f) Pdgfrβ-CreRha-Tom line had numerous pericytes labeled without tamoxifen treatment. (g) After tamoxifen ip, all mural cells in Pdgfrβ-CreRha-Tom line, were positive for TdTomato; in addition majority of Muller cells were labeled (magnified area). (h) After eye tamoxifen drops, all mural cells were labeled, many Muller cells were also positive. Scale bars: 100 um in a-c and f-h, 25 um in d-e.

When we assessed tamoxifen ip induced Cre translocation into the nucleus, the strongest Cre expression was found in pericytes (Figure 3d). After tamoxifen eye drops, we could only detect Cre in pericytes but not SMA cells (Figure 3e). Thus, in Pdgfrb-CreCsln line, activation of Cre in all contractile cells or in pericytes alone reflects the activity of the PDGFRB promoter, known to be more active in pericytes than in smooth muscle cells [25]. In the Pdgfrb-CreRha mouse line, the Cre activation pattern was different (Figure 3f-h). Without tamoxifen treatment, many pericytes expressed TdTomato (Figure 3f) suggesting some “leak” of Cre. Upon both tamoxifen ip or eye drop treatment, all contractile cells were targeted (Figure 3g,h). In addition, the vast majority of Muller cells were also brightly labeled.

### Optimization of pericyte targeting in the retina and brain

The unique specificity of Cre activation in retinal pericytes after tamoxifen eye drops vs ip treatment warrants further investigation as it presents an opportunity to selectively target specific vascular regions for insight in vascular biology and treatment. For this, the animals were treated either with tamoxifen ip, tamoxifen eye drops, or oil ip (Figure 4). A coronal section through the hippocampus was chosen for brain evaluation (Figure 4a). After ip tamoxifen, all major blood vessels and capillaries were targeted in the brain with no positive glia detected (Figure 4a and insert). In the retina of the same mouse, all mural cells were also labeled with no endothelial cells labeled (Figure 4b). Muller cells were weakly positive for TdTomato. Surprisingly, very low numbers of brain pericytes were targeted by tamoxifen eye-drop application (Figure 4C and insert), with no labeling of major blood vessels. The density of targeted pericytes was much lower than in mice treated with tamoxifen injection and just slightly higher than in the animals treated with oil alone (Figure 4e and insert). However, in the retina following eye-drops tamoxifen, the majority of pericytes were labeled and the uninterrupted net of the capillaries could be seen (Figure 4d). TdTomato was found exclusively in mural cells, with the highest expression levels being in pericytes, and with a strong correlation to the high density of PDGFRb (green). No Muller cells were labeled in this line. As expected, oil-treated animals revealed very few pericytes in both the brain (Figure 4e) and retina (Figure 4f).

**Figure 4.**
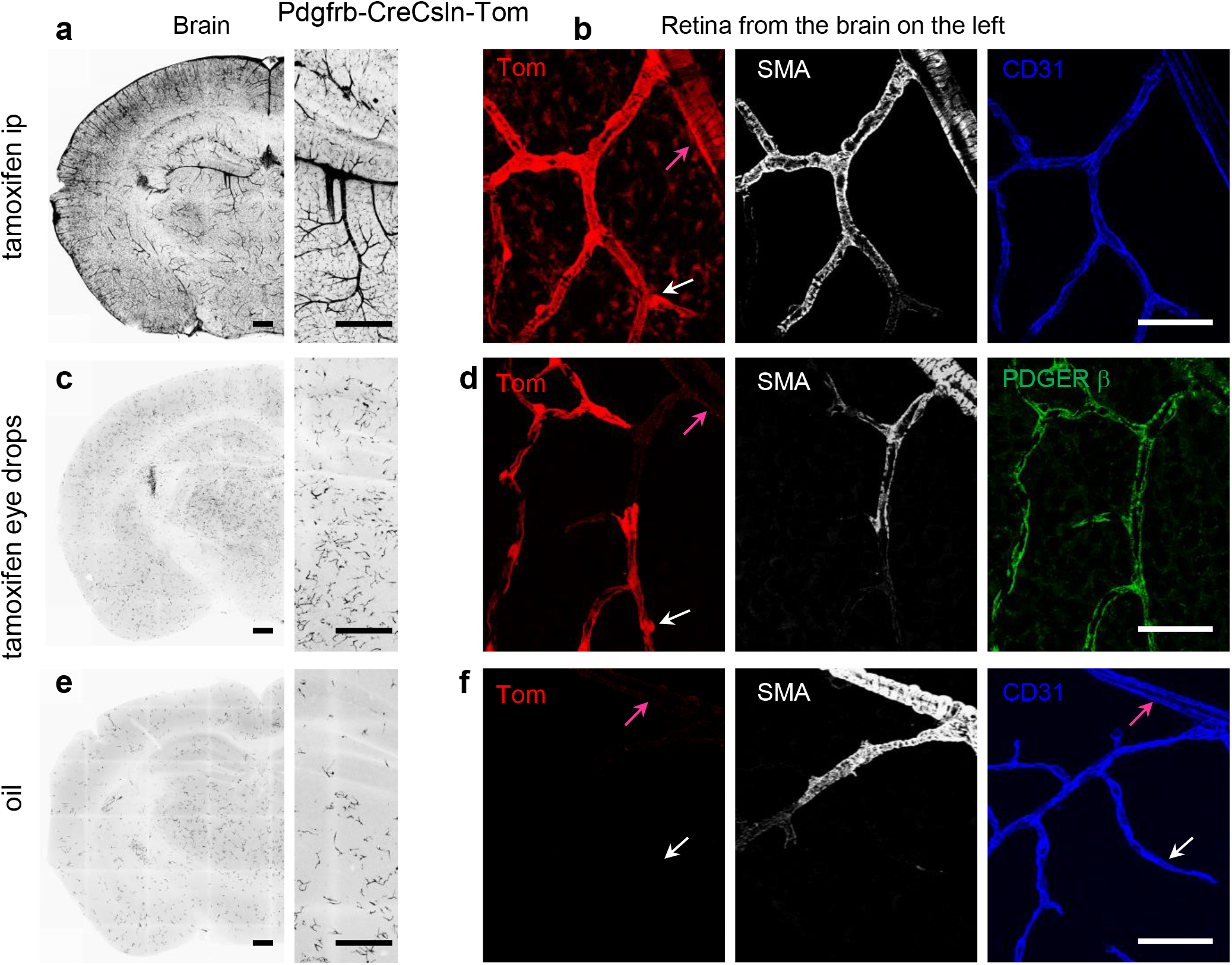
Controlling activation pattern of CreERT2 via tamoxifen administration in B6.Cg-Pdgfrbtm1.1 (cre/ERT2)Csln/J line. (a) In the brain of tamoxifen ip treated Pdgfrβ-CreCsln-Tom all mural cells expressed TdTomato; no glial labeling was visible. (b) In the retina of the mouse from (a), TdTomato-labeling was in SMA cells (magenta arrow) and pericytes (white arrow), as well as many Muller cells. No endothelial cells (blue) were labeled. (c) In the brain of Pdgfrβ-CreCsln-Tom treated with tamoxifen eye drops, TdTomato was found predominantly in the individual separate pericytes in the capillaries but not in large arteries and veins. (d) In the retina of the mouse from (b), TdTomato-labeling was colocalized with Pdgfrβ immunolabeling (green) suggesting selective Cre activity in the majority of pericytes (white arrow) but not in SMA cells (magenta arrow). (e) In the brain of oil ip treated Pdgfrβ-CreCsln-Tom, sparse TdTomato labeling was detected in pericytes. (f) In the retina of the mouse from (e), there was no TdTomato-positive cells neither among SMA cells (magenta arrow) nor pericytes (white arrow). Endothelial cell immunolabeling for CD31 was used to visualize capillaries (blue). Scale bars: 400 um in a, c, and e; 25 um in b, d, and f.

Thus, tamoxifen eye-drop application was a reproducible method of Cre activation for the majority of retinal pericytes yet much less for brain pericytes. Therefore, the organ specific tamoxifen application could be used for retina specific pericyte targeting.

### Leveraging selective vascular cell targeting using optogenetic stimulation for manipulation of retinal blood flow

Next, we evaluated the ability of the Cre system to produce functional changes in mural cells. For this we choose the Pdgfrb-CreCsln mouse line and crossed it with a Halorhodopsin reporter mouse (B6;129S-Gt(ROSA)26Sortm39(CAG-hop/EYFP)Hze/J). Halorhodopsin is a membrane bound chloride pump which induces hyperpolarisation in response to green/yellow light (peak wavelength sensitivity at 570 nm). The offspring were heterozygous for both Cre and Halo, and were treated with tamoxifen eye drops (Figure 5a). Similar to the Pdgfrb-CreCsln-Tom reporter line, Halorhodopsin was selectively expressed in the majority of pericytes in the Pdgfrb-CreCsln-Halo line (Figure 5b). However, in contrast to TdTomato expressed in the cytoplasm, the Halo-YFP construct was localized at the membrane of pericytes, consistently with its function as transmembrane ionic channel. No endothelial or glia cells expressed Halorhodopsin. Tamoxifen and oil treated mice were used for assessment of retinal blood flow under a two-photon microscope (Figure 5c). Mice were lightly anesthetized and iv injected with the fluorescent contrast probe Sulforhodamine 101. The 10x objective enabled precise visualisation of each of the three vascular layers when changing the focal plane. Blood flow was measured in individual capillaries in each layer, in ROIs of 128 by 128 pixels (55 by 55 um), (highlighted in red, Figure 5) before and after activation of Halorhodopsin. Baseline, and response recording epochs lasted for 30 s, and intercalated by a 20 s light-stimulation of green light (peak wavelength 565 nm, power 900 mW).

**Figure 5.**
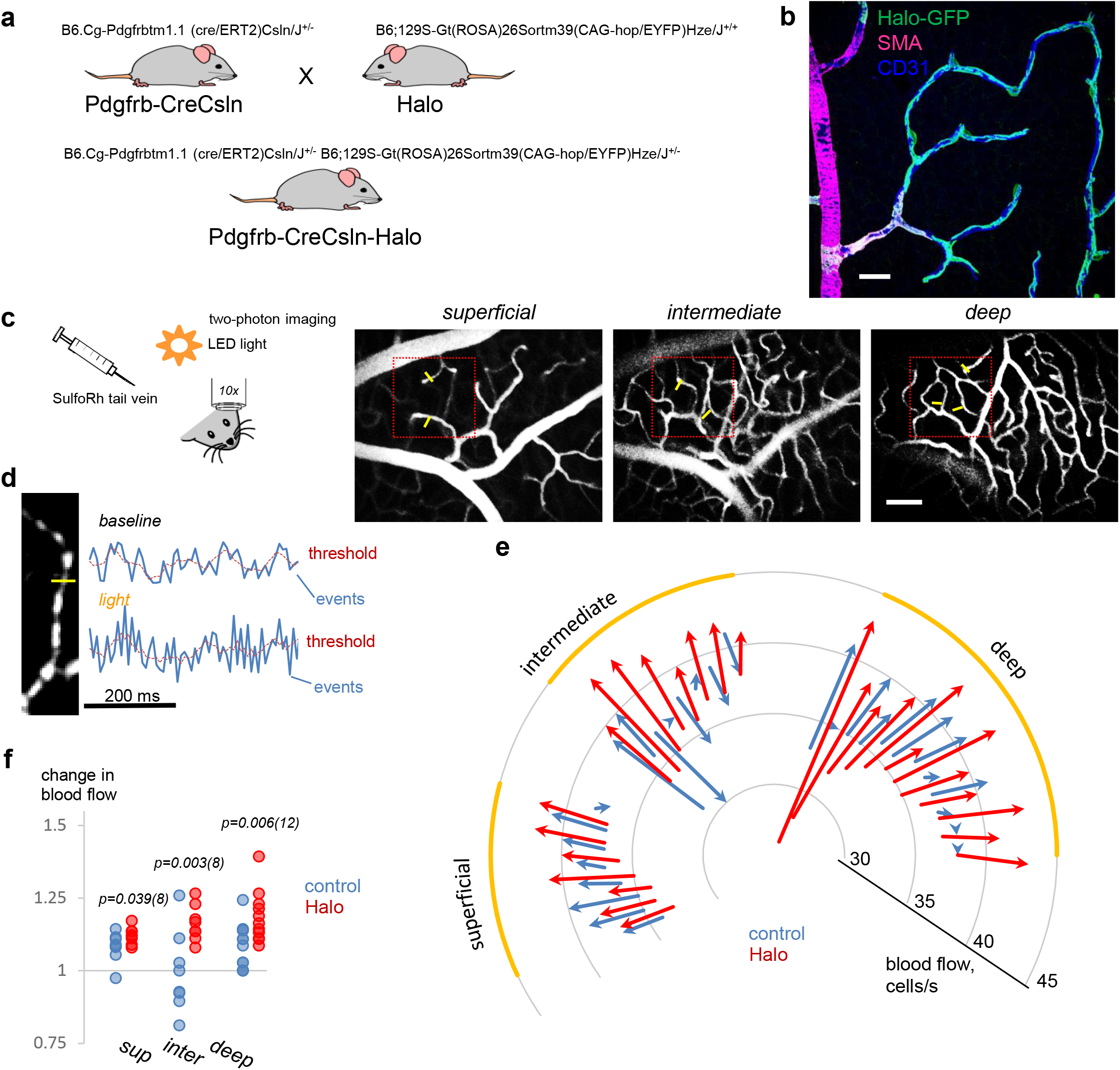
Targeted expression of Halorhodopsin in retinal pericytes for optogenetic boost of capillary blood flow. (a) Creation on Pdgfrb-CreCsln-Halo mouse. (b) Predominant expression of Halorhodopsin by contractile cells in microvascular regions. Retinal wholemount from B6.Cg-Pdgfrbtm1.1 (cre/ERT2)Csln/J^+/−^ and B6;129S-Gt(ROSA)26Sortm39(CAG-hop/EYFP)Hze/J^+/−^ mice expressing Halorhodopsin after Cre-induction via tamoxifen eye drops. Scale bar 25 um. (c) In vivo imaging of retinal vasculature and blood flow after tail vein infusion of sulforhodamine101. Three vascular layers are clearly distinguishable. The actual area of blood flow with fast scan is outlined by the red square. Yellow bars indicate cross-sections through capillaries where blood flow was measured. Scale bar 25 um. (d) Calculation of blood flow was made by analyzing time-lapse images of individual capillaries. Blood cells appear dark relatively to sulforhodamine101-labeled plasma. At the capillary cross-section marked by yellow line, the fluorescence intensity profile was created (blue line on the graph) and passing blood cell events were registered as peaks falling below the threshold. (e) Graphic representation of capillary blood flow before and after light stimulation at each capillary sample across three distinct vascular layers in halo and control animals. A vector indicates changes of blood flow in an individual capillary at the baseline (start) and after light stimulation (end). Four eyes per group. (f) Optogenetic stimulation produced a robust increase in capillary blood flow. Ratios of capillary blood flow after light stimulation to the baseline in *wt* (blue) and Halorhodopsin-expressing retinas (red) were significantly different. Four eyes per group. Numbers of measured capillaries are below data points; TTEST significances are above.

For capillary blood flow assessment, a fluorescent profile was measured across individual capillaries (Figure 5d) and all peaks above the threshold line were counted as passing blood cells (Methods). At baseline, oil and tamoxifen-treated mice had similar blood flows in the superficial (control, 36 ± 1; tamoxifen, 36 ± 1; p = 0.22, TTEST, N=4), intermediate (control, 37 ± 3; tamoxifen, 36 ± 1; p = 0.14), and deep (control, 36 ± 1; tamoxifen, 34 ± 4; p = 0.03, TTEST, N=4) layers (Figure 5e). After Halorhodopsin activation, the blood flow increased significantly in intermediate (control, 37 ± 3; tamoxifen, 42 ± 1; TTEST< 0.001) and deep (control, 39 ± 2; tamoxifen, 41 ± 2; p = 0.02 TTEST; N=4) layers but not in the superficial (control, 40 ± 2; tamoxifen, 36 ± 1; p = 0.2, TTEST, N=4). Interestingly, in the intermediate layer there was an increase of blood flow in the majority of the capillaries in tamoxifen-treated mice but not in the oil-treated mice. Finally, we compared ratios of blood flow increase after stimulation relative to the baseline (Figure 5f). Again, the ratios were significantly different in the superficial (control, 1.08 ± 0.05; tamoxifen, 1.12 ± 0.03; p = 0.039, TTEST, N=4), intermediate (control, 0.99 ± 0.14; tamoxifen, 1.16 ± 0.06; TTEST = 0.004, N=4), and deep layers (control, 1.08 ± 0.08; tamoxifen, 1.22 ± 0.16; p = 0.006; TTEST, N=4) with higher blood flow increase in tamoxifen-treated mice. The expression of Halorhodopsin was confirmed in each animal after the *in vivo* experiment. Indeed, Halorhodopsin was associated with the EYFP reporter tag, which can be observed by fluorescence in the explanted retina. Using optogenetic tools and applying them specifically to modulate pericyte activity, we were able to transiently increase capillary blood flow in the retina of a living mouse.

## Discussion

The retina is one of the most metabolically demanding organs in the body and has been extensively used as a model to study CNS angiogenesis and blood flow. Here, using three commercially available mural cell specific CreERT2 lines, we found that targeting mural cells in the retina is distinct from other parts of the CNS. Specifically, tamoxifen application sets either ubiquitous targeting of all mural cells or pericytes alone. To demonstrate the utility of the pericyte-specific Cre-lines for vascular research, we evaluated the morphology of a newly discovered vascular sphincter cell and boosted capillary blood flow by optogenetically modulating pericyte physiology *in vivo*. Below, we provide a comparative analysis of pericyte targeting approaches and discuss how they can be used for basic and translational studies.

### Which line to choose? Comparative analysis of mural-specific mouse lines in the retina

The three investigated CreERT2mouse lines for inducible targeting of mural cells exhibited different baseline, selectivity, and density of tamoxifen-activated mural cells (Table 2).

**Table 2.** Mouse lines evaluated for targeting vascular mural cells in the retina.

| Name of the line | Background activation | Tamoxifen ip | Tamoxifen eye |
| --- | --- | --- | --- |
| Pdgfrb-CreRha | many pericytes | all mural cells and many Muller cells | all mural cells and many Muller cells |
| Pdgfrb-CreCsln | very few pericytes | all mural cells and some Muller cells | predominantly pericytes and a few SMA cells |
| Cspg4-Cre | very few random type mural cells | sparse random type mural cells | sparse random type mural cells |

The main advantages of the CreERT2 system is that Cre-mediated recombination of floxed target sites can be induced at a specific time through tamoxifen administration [26,27]. Cre leakiness is a known concern, and may be due to incomplete cre sequestering in the cytoplasm when a large amount of cre protein is produced [28,29]. Indeed, in the Pdgfrb-CreRha-Tom line, numerous pericytes were labeled prior to tamoxifen treatment. The reporter lines are known to exhibit different recombination thresholds [29,30]. We used a highly sensitive Ai14 reporter [8], highlighting that Cre in the Pdgfrb-CreRha which was “leaky” in combination with Ai14 may have no background activity with a different reporter line [31,32]. Thus, for each CreERT2/loxP system the inclusion of oil-treated controls that carry both the CreERT2 and the reporter allele, is a necessity. Our data also show that the widely used Ai14 reporter line has some background TdTomato expression in Muller glial cells in the retina.

The other important characteristic is the specificity of Cre expression in individual mural cell types. All three lines, when induced by tamoxifen via ip injections, targeted all types of mural cells. In the Pdgfrb-CreRha-Tom line, additional labeling was found in Muller cells. The selectivity of Cre activation, especially at low tamoxifen concentration, was strongly defined by the activity of its promoter. In the adult mice, the Cspg4 promoter is active in all mural cells [33,34], while the Pdgfrb promoter is predominantly active in pericytes and transitional cells within the precapillary regions and less in arterial SMA-positive cells [35]. In our study, PDGFRb immunolabeling was strongly associated with pericytes and less with the SMA-positive cells. The degree and the pattern of Cre activation may also depend on the route or concentration of tamoxifen administration. Interestingly, we found that in the Pdgfrb-CreCsln line, tamoxifen eye drops induced Cre in the vast majority of retinal pericytes and to a significantly lesser degree in the brain of the same animal (Figure 4). The high-specificity and simplicity of our approach opens the opportunity to selectively manipulate pericytes without affecting SMA cells. Furthermore, we demonstrate that focal eye drop tamoxifen application restricts Cre activation within the retina, minimizing undesired off-target effects.

### Mural cell-specific CreERT2 lines for basic research

The CreERT2 system, in combination with floxed genes allows visualization of cells, overexpression or suppression of their genes, cell ablation, inhibition, and excitation [36,37]. By combining mural cell promoter lines with fluorescent reporter lines, we enabled the visualization of individual mural cells for detailed morphological characterization. First, using the Cspg4-Cre-Tom line, we anatomically characterized sphincter pericytes. In the brain, they control blood inflow from an artery into a capillary branch [21,38]. We found that this is a single cell fully encircling the opening into the branching vessel; the cell had relatively low SMA expression and was heavily insulated by the basement membrane. Insulation of the sphincter suggests signal separation of the major artery from the capillary branch. This is supported by physiological studies that demonstrated a cutoff point for vasomotor signaling between artery and capillary branch through electrical stimulation, calcium imaging and neurobiotin tracing [23,22]. Second, sparse labeling of individual pericytes enabled us to reveal their structural polarity. This structural asymmetry correlated with the direction of blood flow. Pericytes wrapped capillaries in the upstream direction of blood flow and had a long narrow tail on the downstream side. It is possible that this anatomical polarity serves functionally distinct regulatory mechanisms needed for proper control of vascular signaling and blood flow [22,39,40].

Studies in pericyte-deficient mice showed that pericytes control blood-brain/ blood-retinal barriers [41]. To dissect the BBB-supporting role of pericytes in development and mature vessels, tamoxifen inducible lines could be crossed with Cre-inducible diphtheria toxin receptor lines [42]. Instead of cell ablation, their genes could be manipulated to determine the molecular players involved in the BBB supporting function of pericytes. For example, Foxf2 inactivation in adult mice results in BBB breakdown, endothelial thickening, and increased trans-endothelial vesicular transport [43].

### Targeted manipulation of pericytes to boost capillary blood flow

There are a number of diseases in the retina and brain that are affected or driven by pericyte pathology, starting with the disruption of physiological processes (microcirculation dysfunction) and progressing onto maladaptive anatomical remodeling (edemas, neovascularization). We sought to target pericytes to enhance capillary blood flow as recent studies identify them as key players in both the sensing and delivery of localized functional hyperemia in the retina [22].

A recent study explored the possibility to elicit light-driven capillary constriction and reduce blood flow *in vivo* by depolarizing brain pericytes with the cation optogenetic channel ChR2 [44]. As we sought to increase local blood flow by relaxing pericyte tone, an optogenetic hyperpolarizing system presented an ideal opportunity. The light-sensitive anion channel halorhodopsin [45] enabled hyperpolarization of pericytes, resulting in a local dilation of capillaries, thus bypassing the highly orchestrated process of functional hyperemia [10]. Indeed, targeting of multiple vascular elements simultaneously on the vascular tree leads to paradoxical effects which cancel each other out [46], warranting the need for highly precise cellular and spatio-temporal targeting with optogenetics. We would like to acknowledge that Halorhodopsin may present a number of limitations as an effective treatment strategy, requiring high photopic irradiance levels for activation, and operating on a millisecond timescale [45]. Possible alternatives include step-function opsins (bi-stable ChR2 mutants designed to stabilize the active chromophore isomer) which present longer inactivation time constants as well as single light-pulse switch-off mechanisms [47]. Animal opsins (type II rhodopsins) require a much lower irradiance for activation, but activate G-protein pathways [48], whose actions remain to be dissected in contractile pericytes. Similarly, designer receptors exclusively activated by designer drugs (DREADDs) can provide acute yet sustained G-protein activation in response to synthetic agonist administration with highly selective cellular targeting [49].

In diabetic neuropathies, blood flow reduction in human patients [50] and animal models [51,52] was established long before pericyte drop-out, neovascularization and BBB breakdown. Targeted enhancement of blood flow with optogenetics or DREADDs could optimize oxygen delivery and avoid development of hypoxia-induced neovascularization and BRB impairments. In retinal degenerative (RD) diseases, blood vessel degeneration was commonly considered to be secondary to neuronal deficits. However, recent studies showed that narrowing of blood vessels and their consecutive degeneration was con-current to neuronal deficits [16,53]. Moreover, the preservation of blood vessels was sufficient to rescue cone photoreceptors in some rodent models of RD [54]. In humans, Retinitis pigmentosa is characterized by an initial reduction in the blood flow beyond changes associated with the degradation of blood vessels [55–57,53,58]. Persistent blood flow through capillaries was shown to be essential for the survival of capillaries during the pruning process [59]. Moreover, pericytes persisted, even on the dead blood vessels as long as they had a point access to a living blood vessel [16]. Their persistence makes them a good target for optogenetic or DREADD support to improve microcirculation in RD.

## Acknowledgments

This work was supported by NIH grants R01-EY026576, R01-EY029796 and NYSDOH-C34457GG (BTS).

